# Discovery of P_1_-P_2_ depeptidized boronic acid based *Plasmodium falciparum* SUB1 inhibitors

**DOI:** 10.64898/2026.10.28.762536

**Authors:** Armands Kazia, Elina Lidumniece, Chrislaine Withers-Martinez, Owain Donnelly, David A. Fidock, Michael J. Blackman, Aigars Jirgensons

**Affiliations:** National Institute of Research and Innovation, Riga, LV-1006, Latvia; Faculty of Natural Sciences and Technology, Riga Technical University, Paula Valdena Str. 3, LV-1048 Riga, Latvia; Malaria Biochemistry Laboratory, The Francis Crick Institute, London NW1 1AT, United Kingdom; Structural Biology Science Technology Platform, The Francis Crick Institute, London NW1 1AT, United Kingdom; Department of Microbiology & Immunology, Columbia University Irving Medical Centre, New York, 10032, NY, USA; Center for Malaria Therapeutics and Resistance, Division of Infectious Diseases, Department of Medicine, Columbia University Irving Medical Centre, New York, 10032, NY, USA; School of Life Sciences, University of Dundee, Dundee, Scotland, DD1 4HN, United Kingdom; Faculty of Infectious and Tropical Diseases, London School of Hygiene & Tropical Medicine, London WC1E 7HT, United Kingdom

**Keywords:** Malaria, *P. falciparum*, serine protease, inhibitors, boronic acids

## Abstract

The *Plasmodium falciparum* subtilisin-like serine protease 1 (PfSUB1) is an essential enzyme required for parasite egress from host red blood cells and represents a promising target for antimalarial drugs. Peptidic boronic acids are potent reported PfSUB1 inhibitors, however their drug-likeness is limited by their pronounced peptidic character. Based on molecular docking studies using peptidic boronic acids, we designed a series of partially depeptidized PfSUB1 inhibitors by replacing the P1–P2 dipeptide fragment with a 3-aminomethylphenylboronic acid core. Structure–activity relationship (SAR) investigations targeting interaction with active-site residue Lys138 revealed that introducing a meta-carboxylate group (compound **3a**) markedly enhanced inhibitory activity, yielding a low-nanomolar PfSUB1 inhibitor (IC_50_= 10.2 nM). Bioisosteric replacement of the carboxylic acid demonstrated that an *N-*acylsulfonamide group (compound **4k**) retained high enzymatic potency (IC_50_= 13.5 nM)). In cellular assays against blood-stage P. falciparum (3D7 strain), the leading depeptidized boronic acids exhibited growth inhibition with values in the low-micromolar range suggesting that further structural modifications are needed to enhance membrane permeability across host and parasite membranes to reach exonemes prior to egress. Overall, this study validates the P1–P2 depeptidization of boronic acid scaffolds and provides key SAR insights for the rational design of non-peptidic PfSUB1-targeted antimalarials.

## 1. Introduction

Malaria is a life-threatening infectious disease caused by *Plasmodium* parasites and transmitted by infected *Anopheles* mosquitoes [1]. It remains a major global health challenge, placing nearly half of the world’s population at risk in tropical and subtropical regions and causing approximately 600,000 deaths annually. Significant efforts have been invested in reducing malaria morbidity and mortality including chemoprophylactic and chemotherapeutic interventions [2]. However, the current antimalarial drug arsenal is increasingly challenged by the spread of drug-resistant *Plasmodium* parasites. To address the growing problem of drug resistance, the most promising strategies involve the development of drugs with novel modes of action capable of disrupting critical stages of the *Plasmodium* life cycle [3-7].

All known *Plasmodium* species posses a single orthologue of subtilisin-like serine protease SUB1, an enzyme essential for merozoite egress from red blood cells (RBC) [8-13]. Shortly before egress, SUB1 is released from exonemes and activates a number of proteins in the parasitophorous vacuole and on the merozoite surface, triggering rupture of both the parasitophorous vacuole membrane (PVM) and the RBC membrane.Disruption of the SUB1 gene has been shown to block merozoite egress during both the liver stage and the subsequent asexual blood stages of infection. The critical role of SUB1 for parasite survival renders this enzyme a promising target for antimalarial drug discovery.

Several SUB1 inhibitors have been identified through either compound library screening or rational design approaches [13-25]. Potent inhibitors have been developed by linking peptidic substructure with reversible covalent serine binding warheads such as β-ketoamides,[14, 24, 25] difluorostatones,[16, 17] boronic acids [21, 22] and irreversible covalent warheads such as diarylphosphonates [23]. So far, peptidic boronic acids **1** are the most potent PfSUB1 inhbitors with low-nanomolar inhbitory potency (Figure 1) [21, 22]. Several compounds (representative compound **1a**, Figure 1) from this series blocked *P. falciparum* egress from infected RBCs and demonstrated submicromolar parasite growth inhibition in cellular assays, but were also found to have off-target effects on the human proteasome. Importantly, several peptidic boronic acids (representative compund **1b**, Figure 1) were shown to exert their parasite growth inhibition via on-target action against SUB1. The selectivity of peptidic boronic acid **1b** for SUB1 *versus* **the** human proteasome (H20S) was achieved at the expense of parasite growth inhibition, with activity in parasite assays reaching low micromolar level.

**Figure 1.**
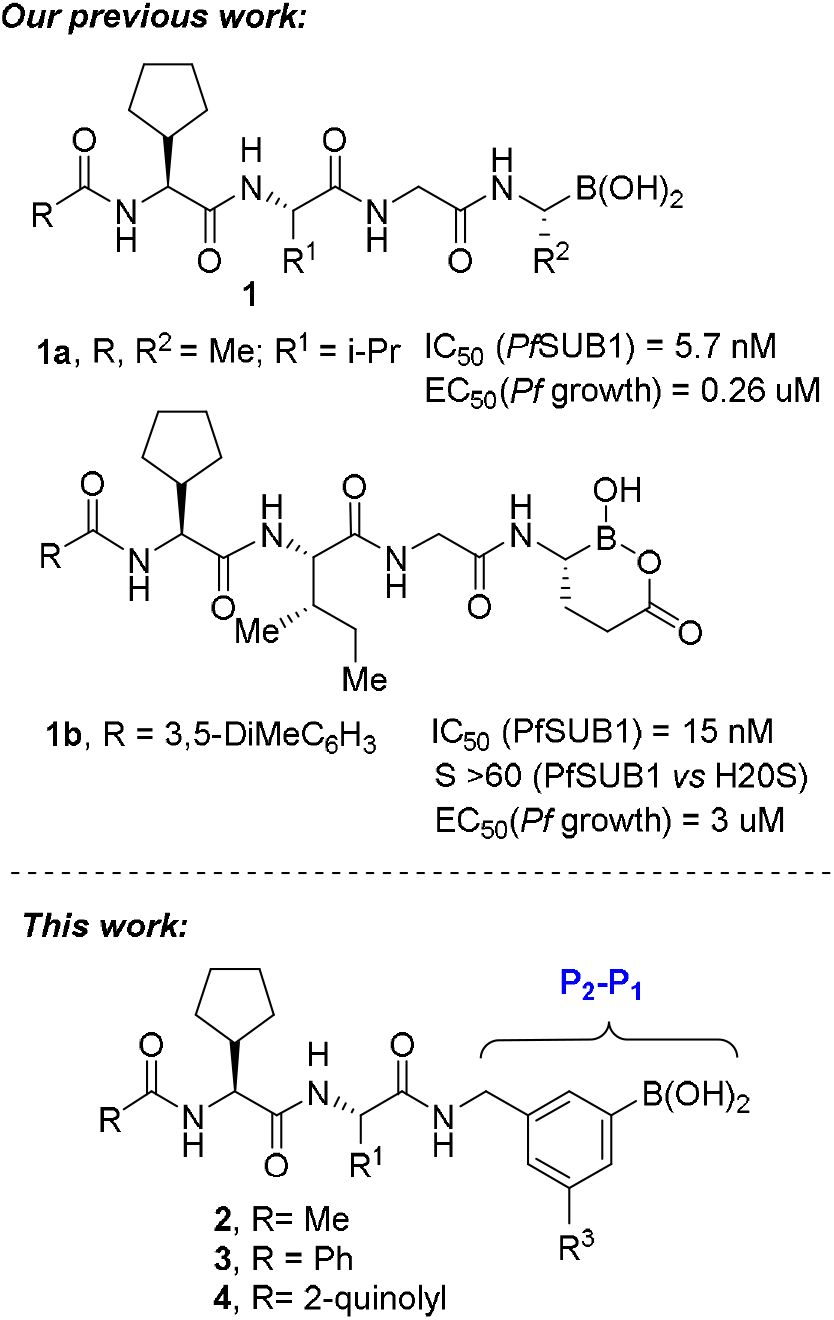
Development of peptidic boronic acids **1** into P_1_-P_2_ depeptidized analogues **2-4**

As a part of our PfSUB1 inhbitor discovery program we aimed to obtain more drug-like inhibitors. One of the avenues explored to achieve this aim was depeptidization of the peptidic boronic acid scaffold, starting with P_1_ – P_2_ peptide segment. Here we report synthesis and structure-activity relationships of partially depeptidized PfSUB1 inhibitors **2**.

## 2. Results and discussion

### Design

A docking model of peptidic boronic acid **1a** bound to PfSUB1 (Figure 2) was used to design depeptidized analogs. Inspection of P2 glycine binding of inhibitor **3a** revealed that the carbonyl group is not involved in hydrogen-bond binding to the enzyme. This indicated the possibility of replacing the P_1_ – P_2_ peptide segment (glycine-alanine) with 3-aminomethylphenylboronic acid (compounds **2-4**, Figure 1). Docking of deptidized analogue **2** revealed good overlap with inhibitor **1a**. The only interaction lost in depeptidized inhibitor binding was the hydrogen bond from NH as donor present in the P_1_ – P_2_ amide linkage of peptidic boronic acid **1a**. Importanly R^3^ substituent in compounds **2-4** pointed to Lys 138. Consequently, this postion was the primary focus for structure-activity relationship investigations.

**Figure 2.**
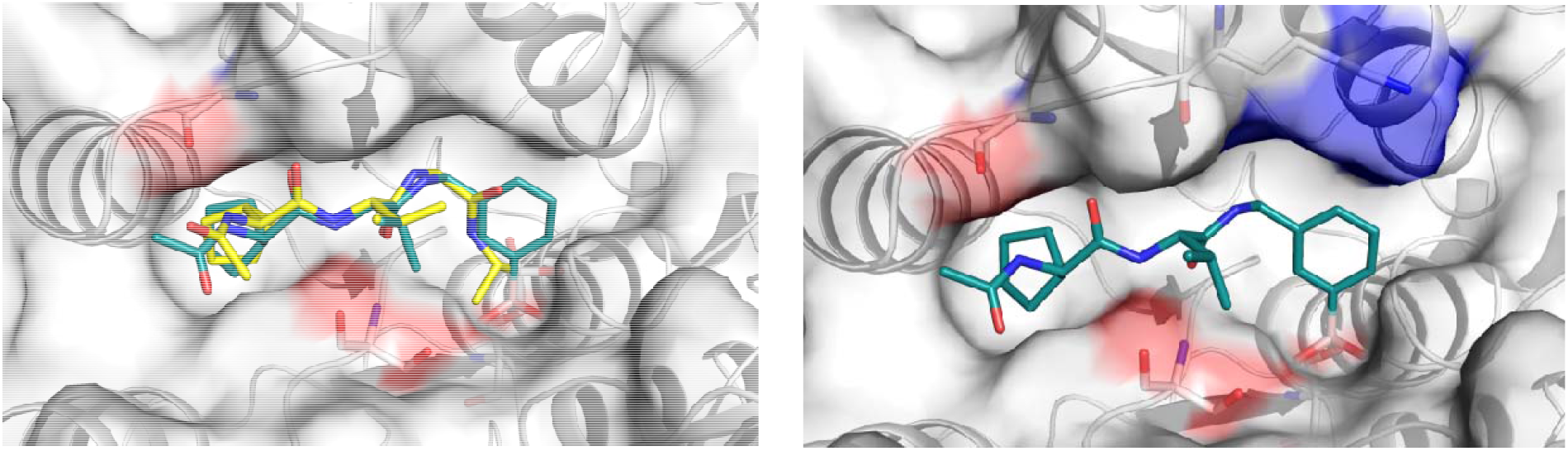
(a) Docking of peptidic boronic acid **1a** overlayed with the docking o**f** 3-aminomethylphenylboronic acid **2** (R= Me, R^3^=H) into the active site of PfSUB1; (b) docking o**f** 3-aminomethylphenylboronic acid **2** (R= Me, R^3^=H) into the active site of PfSUB1with Lys 138 highlighted in blue.

### Synthesis

Peptidic building blocks **8a-8c** for the synthesis of inhibitors **2-4** were prepared starting from *N-*Boc-cyclopentylglycine (**5**) (Scheme 1). This was coupled with methyl esters of L-valine and L-isoleucine to give dipeptides **6a**,**6b**. *N*-Deprotection was followed by *N-*acylation to give acetyl-benzoyl-, and quinoline-2-carboxy-*N-*capped dipeptide esters **7a-7c**. Hydrolysis of intermediates **7a-7c** provided *N-*capped dipeptides **8a-8c**.

**Scheme 1.**
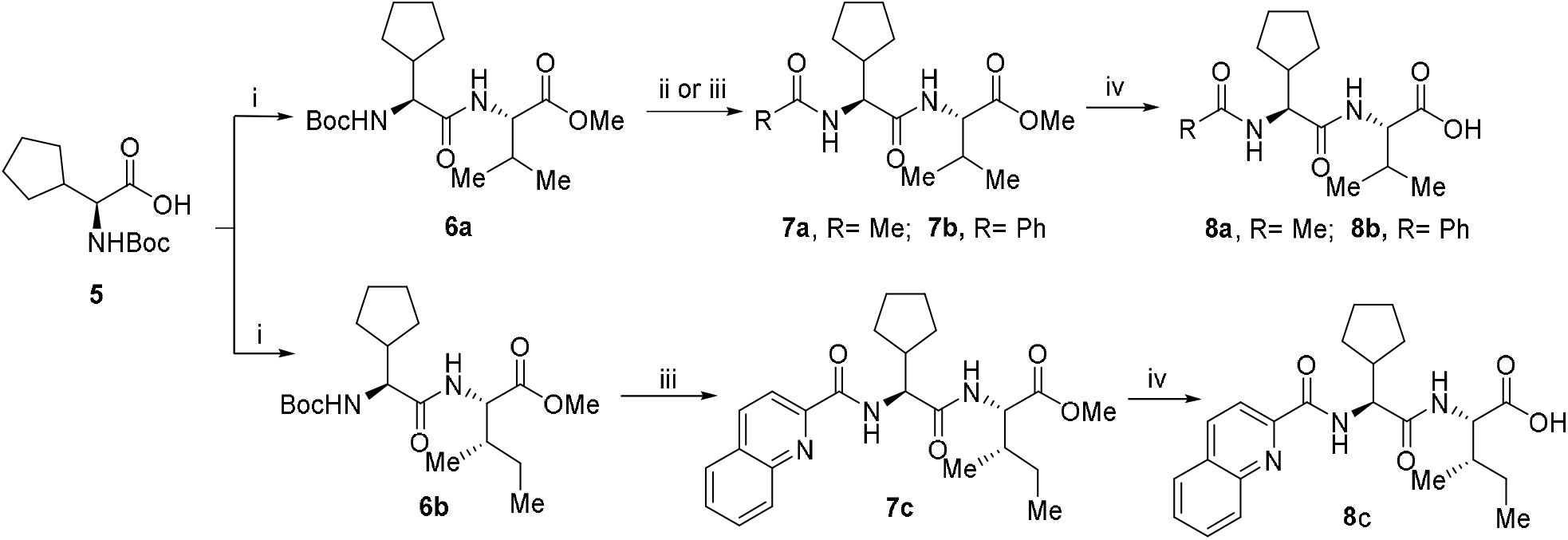
Synthesis of peptidic building blocks **8a-8c. Reagents and conditions:** (i) H**-**L-Val-OMe·HCl or H-L-Ileu-OMe·HCl, EDC·HCl, HOBt, DIPEA, CH_2_Cl_2_ or CHCl_3_, r.t.; (ii) 4 M HCl, CH_2_Cl_2_ then Ac_2_O, DIPEA; (iii) 4 M HCl, CHCl_3_, r.t. then benzoic acid or quinaldic acid, EDC·HCl, HOBt, DIPEA, CHCl_3_, r.t. (iv) LiOH, THF, H_2_O, r.t.

Aminomethylbenzeneboronic acid building blocks **14a-14e** were prepared starting from substituted bromobenzenes **9a-9e** according to Scheme 2 (See Supporting Information for the synthesis of bromobenezenes **9**). Palladium catalyzed coupling of bromides **9a, 9b** with pinacol diborane provided borane pinacol esters which were transformed to more stable (+)-pinanediol esters **10a, 10b**. The same sequence of transformations was used to obtain (+)-pinanediol esters **11c-11e** from bromides **9c-9e**. Formyl and ester groups in borane esters **10a, 10b** were reduced to give hydroxymethyl derivatives **11a, 11b**. These were transformed to bromomethyl derivatives **12a, 12b** by Appel reaction. In turn, O-TBS protected analogues **11c-11e** were first deprotected by exposure to a catalytic amount of Bi(OTf)_3_ and then transformed to iodomethyl derivatives **12c-12e** by the modified Appel reaction using diiodoethane and triphenylphosphine. The resulting halomethyl derivatives **12a-12e** were used to alkylate potassium phthalimide for C-N bond formation. Cleavage of phthalimide with hydrazine in intermediates **13a-13e** provided the desired building blocks **14a-14e**.

**Scheme 2.**
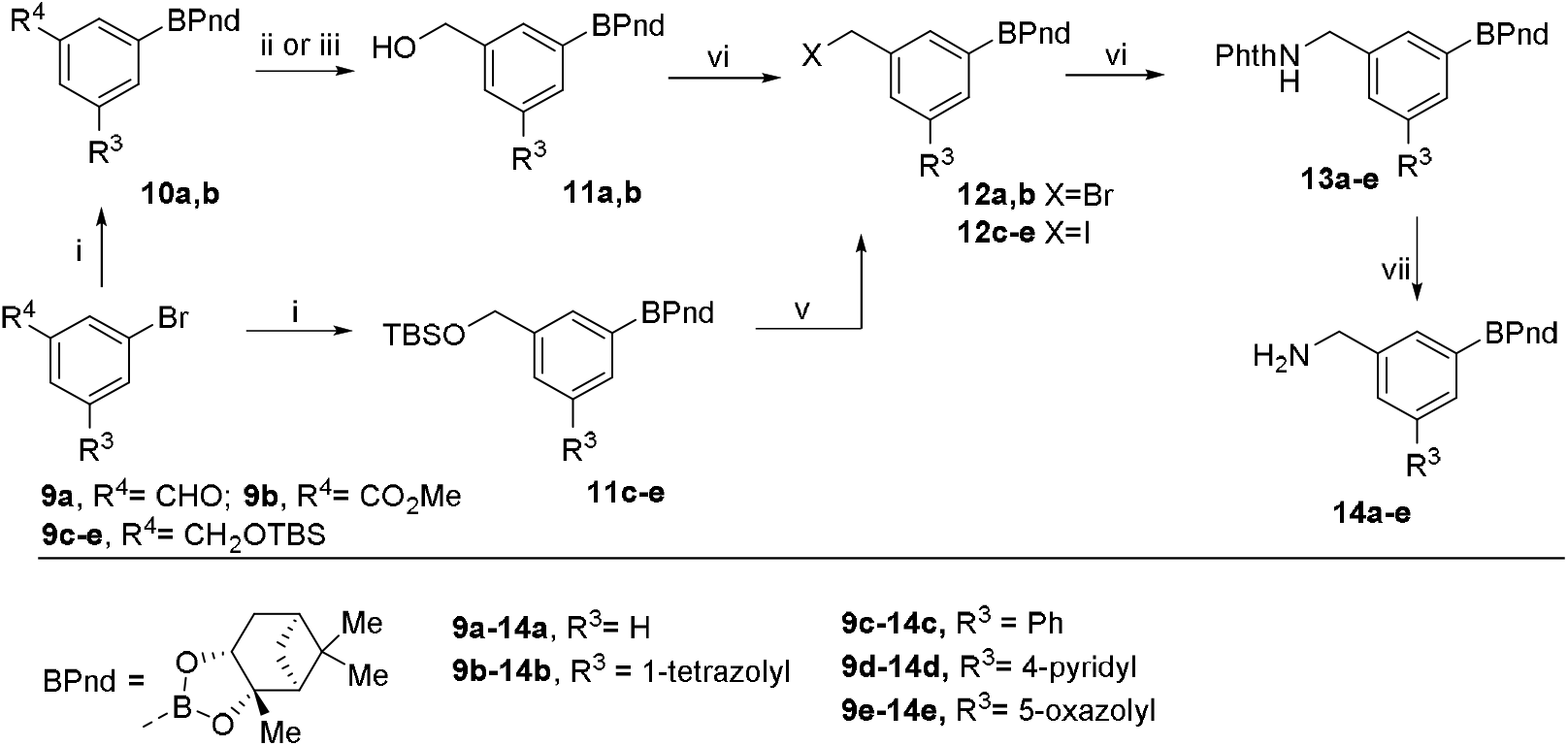
Synthesis of aminomethylbenzeneboronic acid building blocks **14a-e Reagents and conditions:** (i) B_2_Pin_2_, KOAc, cat. Pd(dppf)Cl_2_, dioxane, 90 °C then (+)- pinanediol, THF, r.t.; (ii) NaBH_4_, MeOH, r.t. for **11a**; (iii) LAH, THF, r.t. for **11b**; (iv) CBr_4_, PPh_3_, THF, r.t.; (v) 2 mol% Bi(OTf)_3_, MeOH, r.t. then I-(CH_2_)_2_-I, PPh_3_, MeCN, r.t.; (vi) PhthNK, DMF, r.t.; (vii) N_2_H_4_, THF, r.t.;

Aminomethylbenzeneboronic acid building blocks **18a-18g** were prepared *via* modified synthesis route (Scheme 3). First, hydroxymethylphenyl bromide derivatives **15a-15f** were converted to *N*-alkylated phthalimide derivatives **16a-16f**. Next, these were subjected to Miyaura borylation reaction with pinacol diborane with subsequent transesterification to (+) pinanediol boronates **17a-17f**. Cleavage of phthalimide resulted in building blocks **18a-f**. Aminomethyl derivative **18g** was prepared from *N-*Boc protected aminomethylphenyl bromide **19** by converting it to (+)-pinanediol boronate **20** followed by the cleavage of the Boc group.

**Scheme 3.**
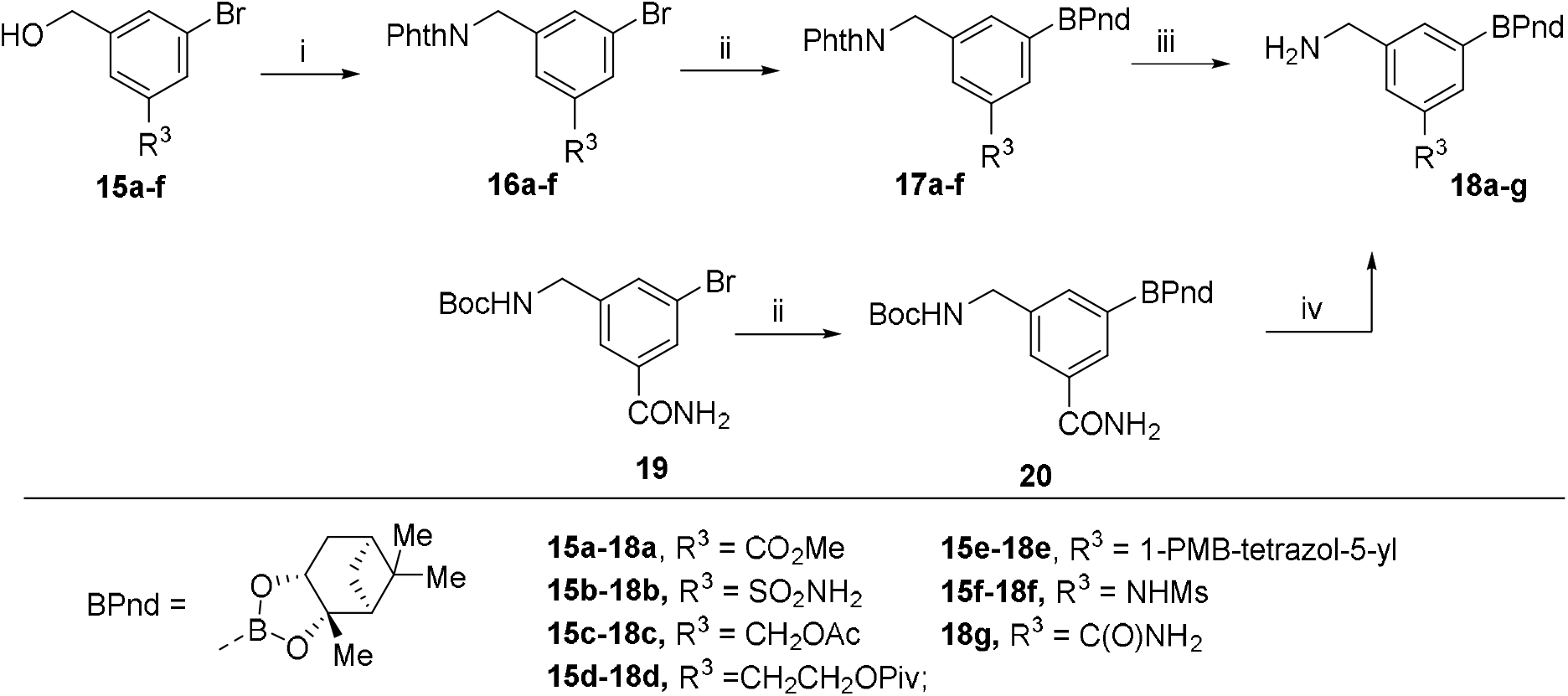
Synthesis of aminomethylbenzeneboronic acid building blocks **18a-g Reagents and conditions:** (i) Phthalimide, DIAD, Ph_3_P, THF, r.t.; (ii) B_2_Pin_2_; KOAc; cat. Pd(dppf)Cl_2_, dioxane, 110 °C; (iii) N_2_H_4_, THF, r.t.; (iv) 4 M HCl in dioxane, DCM, r.t.

Building blocks **8, 14**, and **18** were used to make inhbitors **2-4** (Scheme 4). Dipeptides **8a-8c** were coupled with aminomethylbenzeneboronic acid esters **14a-14e** and **18a-18g** to give intermediates **21, 22a-d**, and **23a-k**. Transesterification of pinanediol boronate precursors **21, 22b, 22c** and **23a, 23c, 23e-23j** with *i*-BuB(OH)_2_ provided free boronic acids **2, 3b, 3c** and **4b, 4c, 4e-4j**, respectively. Hydrolysis of ester functionality in the substitiuent R^3^ of intermediates **22a, 22d, 22e, 23a** followed by transesterification of the boronate ester provided target compunds **3a, 3d, 3e, 4a**. Acid promoted cleavage of PMB group at triazole in intermediate **23d**, acylation and nitrozation of sulfonamide in intermediate **23k** was followed by transesterification of boronate ester to give target compunds **4d, 4k**, and **4l**, respectively.

**Scheme 4.**
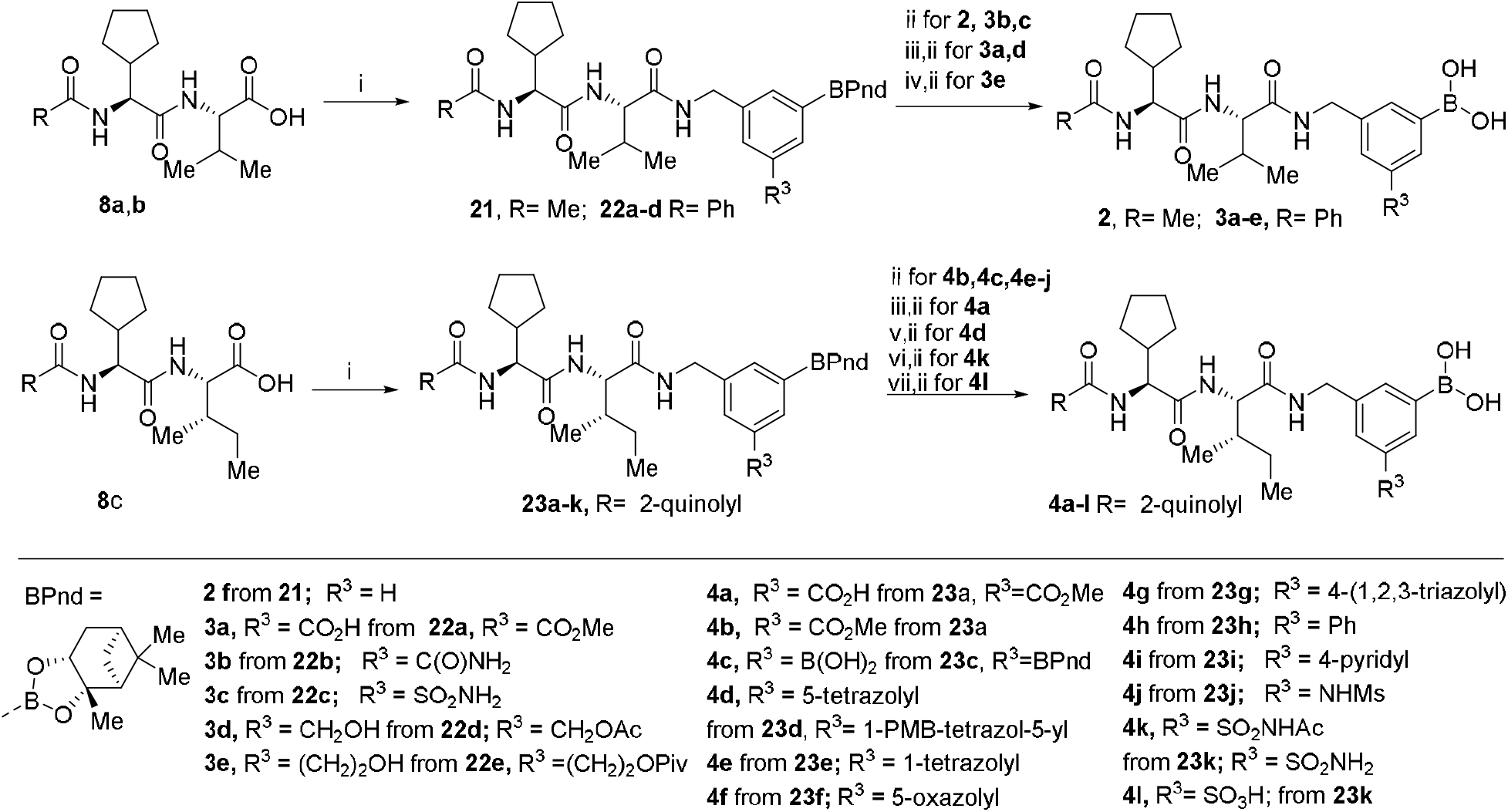
Synthesis scheme of inhibitors **2-4** from building blocks **8a-8c, 14a-14e** and **18a-18g** Reagents and conditions: (i) HATU, DIPEA, DMF, -20 °C to 0 °C; (ii) *i*-BuB(OH)_2_, 1 M aq. HCl, MeOH, *n*-Hex, r.t.; (iii) LiOH, H_2_O, THF; (iv) TBAOH, dioxane water, r.t.; (v) TFA, aniosole, r.t.; (vi) Ac_2_O, DIPEA, DCM. r.t.; (vii) NaNO_2_, TFA, r.t.

An analogue of the peptide-boronic acid conjugate was made with the carboxyl group shifted in ortho-position to boronic acid (compound **28**, Scheme 5). The synthesis started with bromobenzene derivative **24** which was subjected to Miyaura borylation with pinacol diborane followed by tranesterifiaction by pinanediol to give boronate **25**. Boc group cleavage in intermediate **25** provided aminomethyl derivative which was coupled to a peptidic building block **8c**. Ester hydrolysis in coupling product **27** was followed by transesterification of boronate to give target boronic acid **28**.

Furan analogue **33** of boronic acid comprising inhibitors was made from 2-formylfuran-5-boronic acid (**29**) (Scheme 6). This was first esterified with diol to give Epin ester **30**. Formyl group in Epin ester **30** was reduced to hydroxymethyl group and the intermediate was subjected to Mitsunobu reaction with phthalimide providing alkylated phthalimide **31**. Phthalimide group was cleaved with hydrazine and the intermediate aminomethyl furan was acylated with a building block **8c** to give conjugate **32**. Transesterification of Epin ester in compound **32** provided the target boronic acid **33**.

**Scheme 5.**
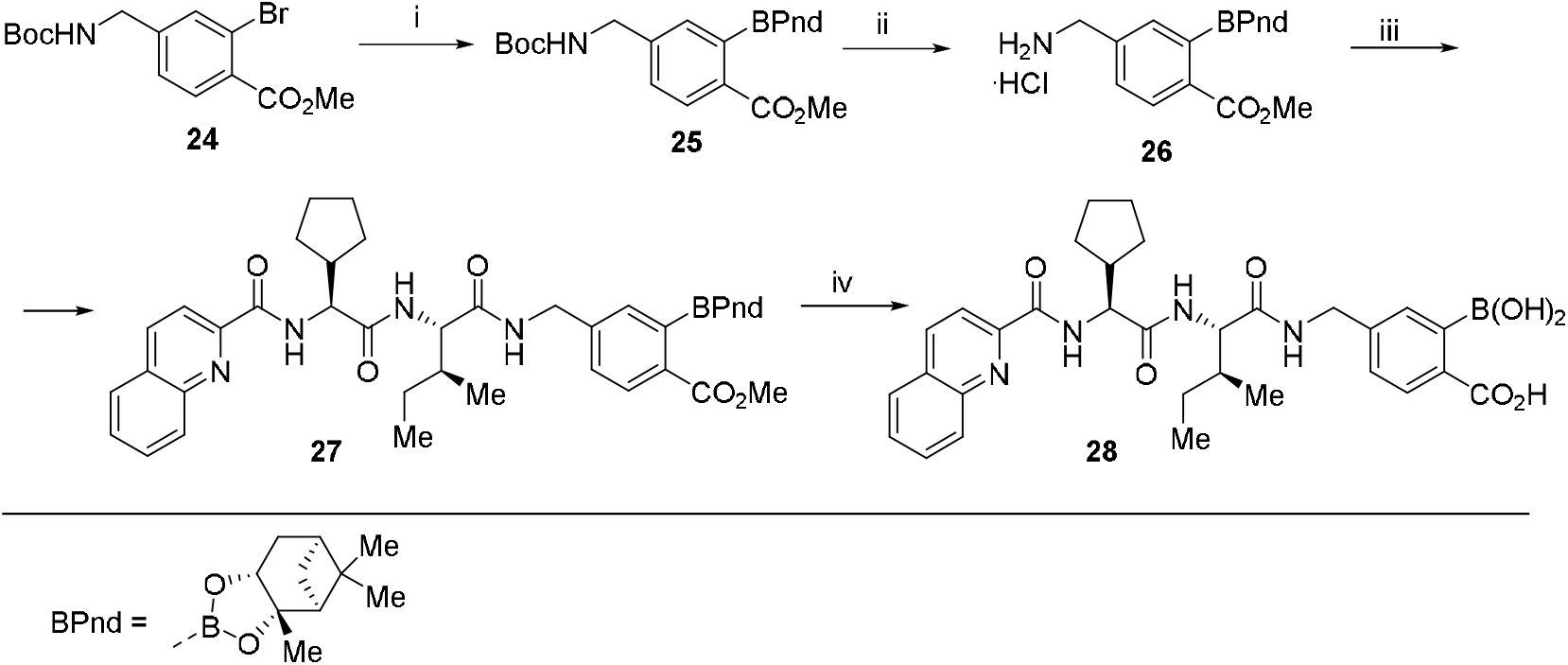
Synthesis of inhibitor **28 Reagents and conditions:** (i) B_2_Pin_2_, KOAc, cat. Pd(dppf)Cl_2_, Dioxane, 90 °C then (+) pinanediol r.t.; (ii) 4 M aq. HCl, CHCl_3_ r.t.; (iii) **8c**, T3P, NMM, CHCl_3_, -15 °C to 0 °C; (iv) LiOH, THF, water, r.t. then *i*-BuB(OH)_2_, 1 M aq HCl, MeOH, *n*-Hex r.t.

**Scheme 6.**
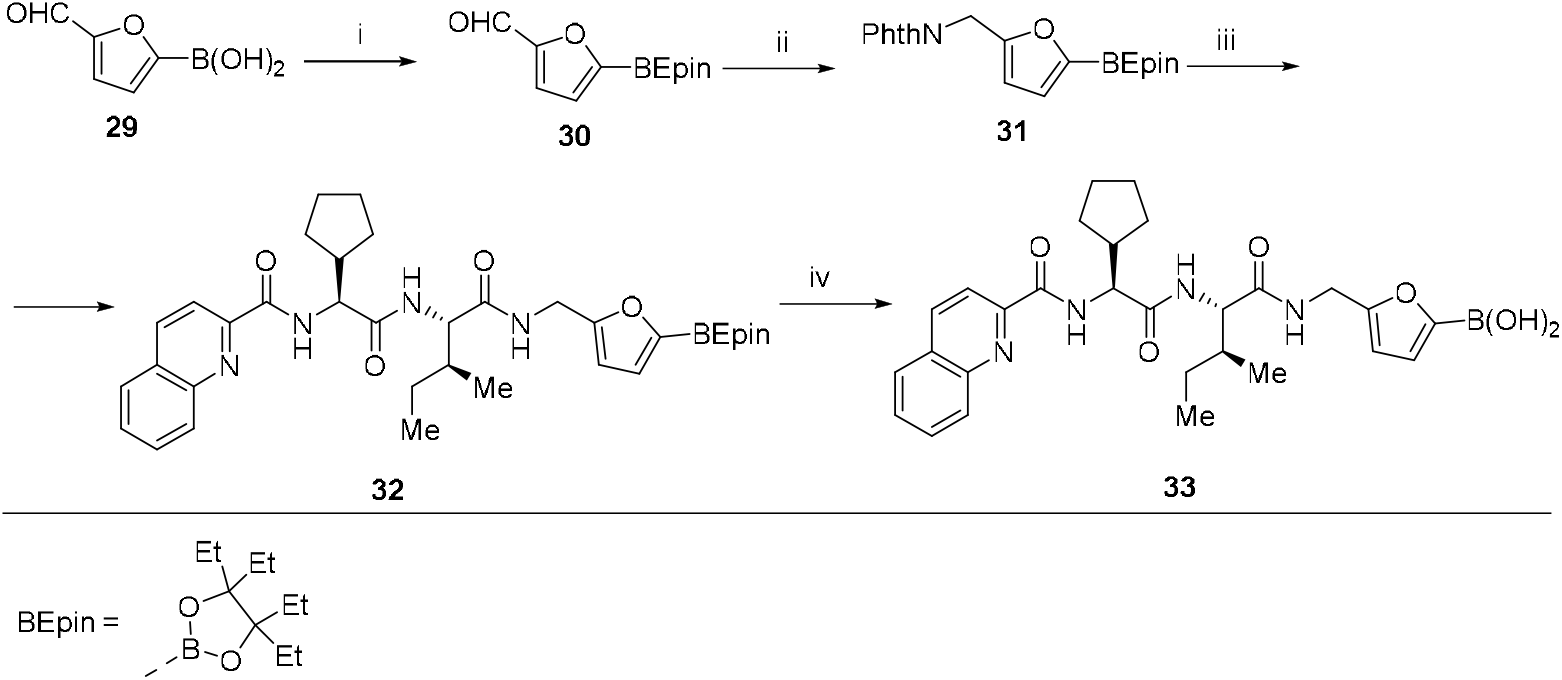
Synthesis of inhibitor **33. Reagents and conditions:** (i) 3,4-diethylhexane-3,4-diol, THF, °C; water, r.t. (ii) NaBH_4_, MeOH, r.t.; (iii) DIAD, Ph_3_P, phtalimide, THF, r.t.; (iv) *i*-BuB(OH)_2_, 1 M aq HCl, MeCN, *n*-Hex r.t.

### Structure-activity relationships (SAR)

Compounds **2, 3a-3e, 4a-g** were tested for their ability to inhibit the PfSUB1 enzyme. The simplest analogue of the series, compound **2** (R^3^= H) showed submicromolar enzyme inhibition potency (Table 1) providing a proof-of concept for benzeneboronic acid-based inhibitors as P_1_□P_2_ depeptidized analogues of peptidic boronic acids **1**. Introduction of carboxylic acid at the meta position to boronic acid (compound **3a**) led to a considerable increase in inhibitory potency. The increase in potency is attributed to a salt bridge formation between the carboxylate of the phenyl group in inhibitor **3a** and Lys138 of PfSUB1 (Figure 2).

**Figure 2.**
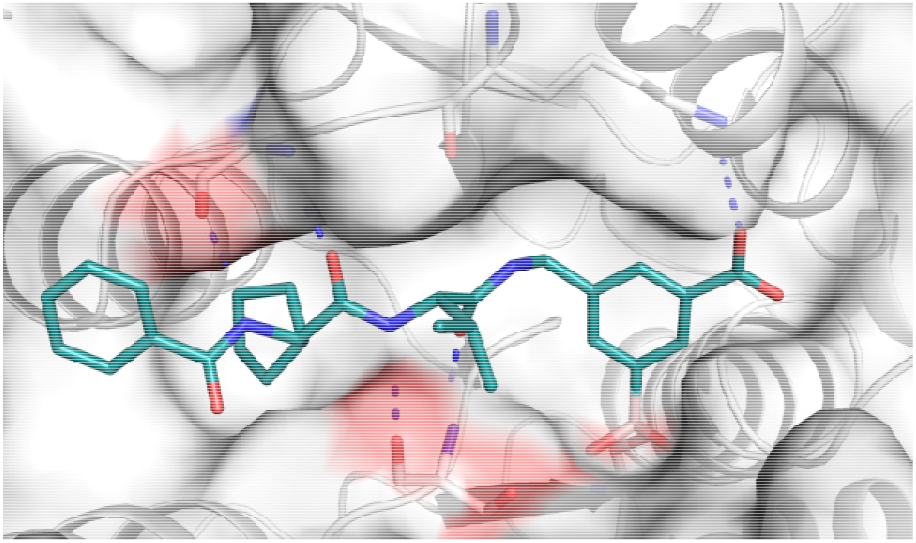
Docking of peptidic boronic acid **3a** into the active site of PfSUB1.

**Table 1.**
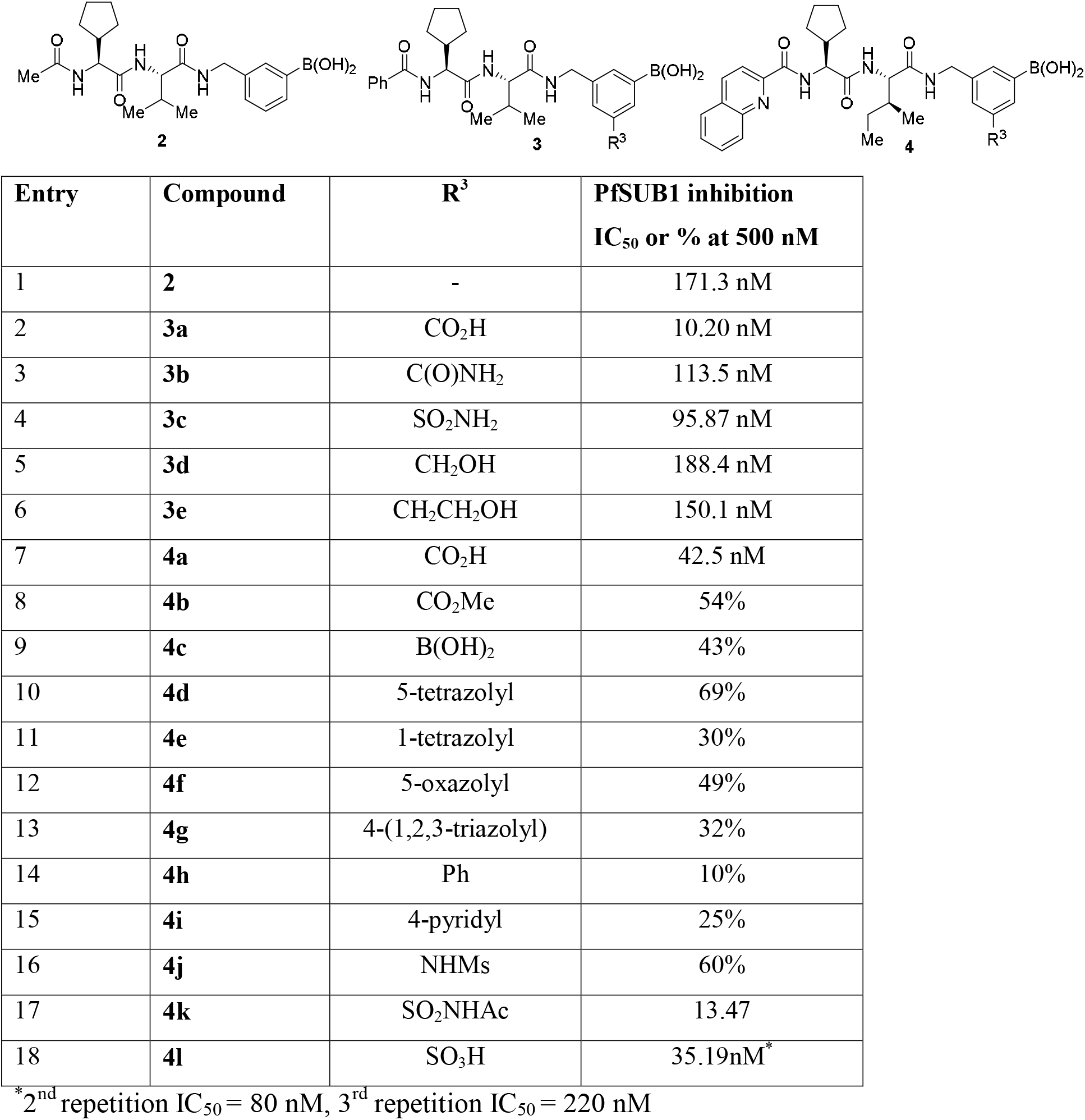
PfSUB1 inhibition potency of 3-aminomethylphenylboronic acid-based inhibitors: SAR of substiuent R^3^.

| **Entry** | **Compound** | **R<sup>3</sup>** | **PfSUB1 inhibition**<br>**IC<sub>50</sub> or % at 500 nM** |
|---|---|---|---|
| 1 | **2** | - | 171.3 nM |
| 2 | **3a** | CO<sub>2</sub>H | 10.20 nM |
| 3 | **3b** | C(O)NH<sub>2</sub> | 113.5 nM |
| 4 | **3c** | SO<sub>2</sub>NH<sub>2</sub> | 95.87 nM |
| 5 | **3d** | CH<sub>2</sub>OH | 188.4 nM |
| 6 | **3e** | CH<sub>2</sub>CH<sub>2</sub>OH | 150.1 nM |
| 7 | **4a** | CO<sub>2</sub>H | 42.5 nM |
| 8 | **4b** | CO<sub>2</sub>Me | 54% |
| 9 | **4c** | B(OH)<sub>2</sub> | 43% |
| 10 | **4d** | 5-tetrazolyl | 69% |
| 11 | **4e** | 1-tetrazolyl | 30% |
| 12 | **4f** | 5-oxazolyl | 49% |
| 13 | **4g** | 4-(1,2,3-triazolyl) | 32% |
| 14 | **4h** | Ph | 10% |
| 15 | **4i** | 4-pyridyl | 25% |
| 16 | **4j** | NHMs | 60% |
| 17 | **4k** | SO<sub>2</sub>NHAc | 13.47 |
| 18 | **4l** | SO<sub>3</sub>H | 35.19nM<sup>*</sup> |

<sup>\*</sup>2<sup>nd</sup> repetition IC<sub>50</sub> = 80 nM, 3<sup>rd</sup> repetition IC<sub>50</sub> = 220 nM

Inhibitors comprising carbocyclic acid bioisosters such as carboxamide, sulfonamide, hydroxymethyl and hydroxy ethyl group showed 10-20-fold lower potency compared to compound **3a**. An analogue of compound **3a** with a quinoline carboxylate end-capping group, compound **4a** was prepared and tested as more lipohilic inhibitor. It was slightly less potent (Table 1, entry 7) compared to the parent compound **3a**. Additionally, for this series several carboxylic acid isosters were introduced, including methyl ester, boronic acid, tetrazole, oxazole and triazole, all of which led to considerably reduced potency. Installation of phenyl, pyridyl or mesylamino group at the meta position relative to the boronic acid abolished PfSUB1 inhibitory potency. *N*-Acylsulfonamide group turned out to be appropriate carboxylic acid bioisostere in compound **4k** leading to improved PfSUB1 inhibitory potency compared to a parent compound. Also, while sulfonic acid comprising compound **4l** was a potent PfSUB1 inhbitor, this compound’s IC_50_ value increased with every repetition, indicating chemical stability issues.

An analogue of inhibitor **4a** with the carboxylic group shifted to ortho position of phenyl boronic acid, compound **28**, turned out to be inactive as a PfSUB1 inhbitor when tested at 500 nM concentration. Also, an analogue of inhibitor **4a** with a furan ring replacing the benzene ring, compound **33**, was inactive as a PfSUB1 inhbitor when tested at 500 nM concentration.

**Figure 2.**
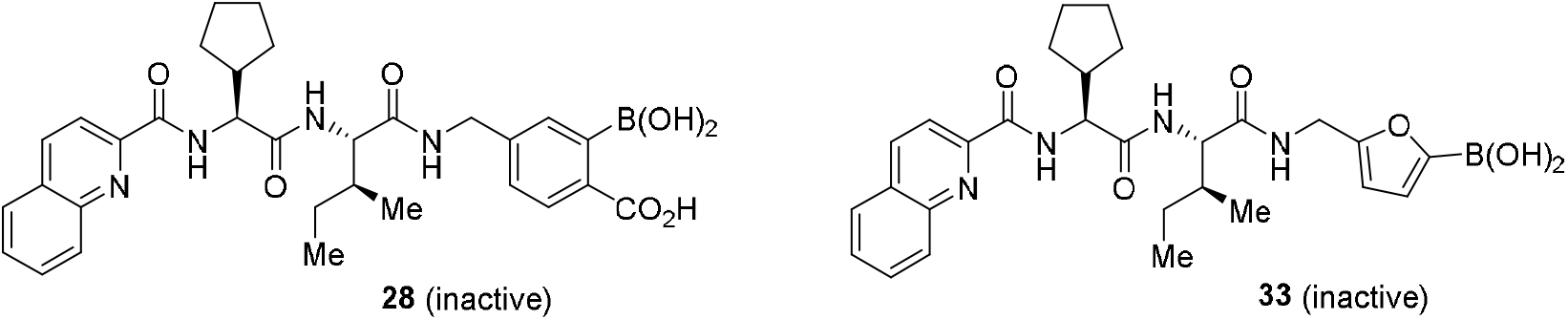
Analogues of inhibitor **4a** with shifted carboxylic group (compound **28**) and scaffold hopping to furan boronic acid (compound **33**).

Selected PfSUB1 inhibitors were tested for their ability to inhibit growth of *P. falciparum* in red blood cells (Table 2). The potency of compounds **3a, 3c, 4e, 4k, 4l** to inhibit the growth of the parasite was found to be in the low micromolar range, similar to compound **1b**, however weaker then for inhibitor **1a** (Table 2). The possible reason of relatively weak potency in a cell-based assay could be due to the poor penetration of inhibitors through the membranes to exonemes where PfSUB1 is stored before the egress, despite the depeptidized character of these inhibitors.

**Table 2.** *P. falciparum* growth inhibition of boronic acids 3a, 3c, 4e, 4k, 4l.

| **Entry** | **Compound** | **R<sup>3</sup>** | **PfSUB1 inhibition IC<sub>50</sub> (nM) or % at 500 nM** | **EC<sub>50</sub> uM<sup>a</sup>** |
|---|---|---|---|---|
| 1 | **3a** | CO<sub>2</sub>H | 10.20 | 16.11 |
| 2 | **3c** | SO<sub>2</sub>NH<sub>2</sub> | 95.87 | 12.06 |
| 3 | **4d** | 5-tetrazolyl | 69% | 25.04 |
| 4 | **4k** | SO<sub>2</sub>NHAc | 13.47 | 18.3 |
| 5 | **4l** | SO<sub>3</sub>H | 35.19 | 9.89 |
| 6 | **1a** | - | 5.7<sup>b</sup> | 0.26<sup>b</sup> |
| 7 | **1b** | - | 15<sup>c</sup> | 3.0<sup>c</sup> |

<sup>a</sup>Parasite growth inhibition was determined using flow cytometry. EC<sub>50</sub> determined by 72 h dose−response assays of the 3D7 strain of *P. falciparum* (N=3, n=2). <sup>b</sup>Data from literature [22]; <sup>c</sup>data from literature [21]

## Summary

In summary, we have developed analogues of peptidic boronic acid inhibitors of SUB1 by replacing P_1_LJP_2_ amino acid part with aminomethylbenzene boronic acid. Introduction of a carboxylic acid into the meta-position of the boronic acid led to considerable (∼10-fold) improvement of SUB1 inbitory potency providing inhibitor **3a** with IC_50_= 10.2 nM. SAR investigation of meta-subsituents of benezene boronic acid revealed that the carboxylic acid can be bioisosterically replaced with *N-*acylsulfonamide providing inhibitor **4k** with IC_50_ = 13.5 nM. Despite their high SUB1 potency, depeptidized inhibitors showed relatively poor (low micromolar) potency to inhibit parasite growth in red blood cell assay. This could be attributed to their poor membrane penetration properties, calling for further depeptidization studies.

## Experimental Section

### Recombinant *P. falciparum* SUB1 inhibition assay

Recombinant purified *P. falciparum* SUB1 (PfSUB1) was produced and purified as previously described [25]. The enzyme was diluted in digestion buffer (25 mM CHAPS, 12 mM CaCl2, 25 mM Tris-HCl, pH 8.2) and dispensed into a white flat-bottomed 96-well fluorescence microtitre plates (Nunc). Test compounds were solubilized in dimethyl sulfoxide (DMSO), serially diluted and added at 2% to the well. The rhodamine-labelled fluorogenic substrate SERA4st1F-6R12 was added at a final concentration of 0.1 μM in a final volume of 100 μl and the rate of hydrolysis was monitored with a Cary Eclipse spectrofluorimeter (Varian, UK) as previously described. Excitation and emission wavelengths used were 552 nm and 580 nm respectively.

### Docking of compounds 1, 2 and 3a into the model of SUB1

Co-folding of compounds **1a, 2** and **3a** and the pfSUB1 sequence (PDB ID: 4LVN) was performed using Boltz-2 (version 2.2.1)[26] and then covalently docked in Schrödinger Maestro (Release 2026-3). The inhibitors were drawn within the Boltz-2 molecule editor using SMILES structures. The PfSUB1-prodomain-NIMP X-ray structure (PDB ID: 4LVN) was converted to a Boltz-2 object for the docking procedure. In this process, hydrogen atoms were added to the SUB1 model, the natural ligand was removed to define the SUB1 active site pocket. The co-folded complexes were transferred to Schrödinger Maestro and a boron-oxygen bond was manually drawn between the boronic acid of compounds **1a, 2**, and **3a** and Ser residue in the active site and minimization was performed. The images were edited in PyMOL (Version 2.6.2 LTS).

### *In vitro* culture maintenance and manipulation of *P. falciparum*

Asexual blood stages of *P. falciparum* strain 3D7 were routinely maintained in human erythrocytes at 3% haematocrit in RPMI 1640 medium supplemented with 25 mM HEPES, sodium bicarbonate (2.1 g/L), gentamicin (10 µL/mL), 50 µM hypoxanthine, 0.5% w/v AlbuMAX II (Thermo Fisher Scientific). Cultures were maintained at 37°C under low oxygen conditions (5% O2 / 5% CO2 / 90% N2 gas mixture), with exchange of culture medium five times per week and fresh erythrocytes added once per week. Human erythrocytes were obtained from anonymized donors through the New York Blood Center and were used within 2 weeks of receipt. Protocol AAAU3761, describing the use of human erythrocytes from anonymized, de-identified donors for *P. falciparum* culture, was approved by the Columbia University Institutional Review Board, who deemed this not to be human subjects research under 45 CFR 46.

### Dose-response assays for assessment of compound EC_50_ in live parasites

The impact of compounds on 3D7 strain P. falciparum growth was assessed using a 72-hour dose response assay. Briefly, a stock concentration (10 mM) of test compounds in DMSO was added by a Tecan D300e digital dispenser to a flat-bottomed, 96-well microtiter plate with twofold serial dilution. Wells were then supplemented with 200 µL per well of sorbitol-synchronised ring-stage 3D7 *P. falciparum* parasite culture at 1% haematocrit, 0.3% parasitaemia. The final concentration of test compound in wells ranged from 50 µM to approximately 195 nM. All drug assays were conducted such that the final DMSO concentration was <0.5%. Assays also included parasite cultures without drug as a negative control and cultures containing 50 µM dihydroartemisinin (a highly lethal antimalarial) as a positive control. Each assay was undertaken on three separate occasions, each time with technical duplicates for each compound (N = 3, n = 2).

Parasitaemia for all wells was measured at 72 hours by flow cytometry (iQue 3, Sartorius) using SYBR Green I and MitoTracker Deep Red FM (Thermo Fisher Scientific) as stains for DNA and cell viability, respectively. Flow counts were analysed by gating the live parasites using FlowJo software (version 10.8.1, FlowJo LLC). Percent parasite growth inhibition (normalized to 100% using the positive control as background) was plotted against log drug concentration and EC50 determined using nonlinear regression using Microsoft Excel (version 16.96.1, Microsoft Corporation). Mean EC50 and standard deviations were calculated from the three biological replicates, each ran in duplicate.

### Chemistry

*General*. All chemicals were used as obtained from commercial sources and all reactions were performed under argon atmosphere in an oven-dried (110 °C) glassware, unless noted otherwise. Anhydrous toluene, Et_2_O, THF, and CH_2_Cl_2_ were obtained by passing commercially available anhydrous solvents through activated alumina columns.

Flash chromatography was carried out using silica gel (230−400 mesh). Thin layer chromatography (TLC) was performed on silica gel and was visualized by UV lamp or staining with KMnO_4_. NMR spectra were recorded on 300, 400 and 600 MHz spectrometers with chemical shift values (δ) in parts per million using residual chloroform, methanol or dimethyl sulfoxide signal as the internal standard. Conversion of starting material was detected with UPLC Waters Acquity, column: Acquity UPLC BEH-C18, 1.7 μm, 2.1 mm × 50 mm, column temperature (30.0±5.0) °C, gradient: 0.01% TFA in water/CH_3_CN 90%/10% – 5%/95%; flow: 0.500 mL/min; time: 8 min; detector: PDA, 220 – 320 nm, SQ detector with an electrospray ion source (ESI/APCI). Gas chromatographic (GC) analysis was performed on Agilent Technologies gas chromatographer with triple-axis detector, heating range 40 – 280 °C, column 30 m × 0.25 mm, 0.25 µm, 7 inch cage. Exact molecular masses (HRMS) were determined on a hybrid quadrupole time-of-flight mass spectrometer equipped with an electrospray ion source. For reversed phase column chromatography Biotage KP-C18-HS SNAP cartridge was used (gradient – water/CH_3_CN).

Synthesis description of compounds are provided in Supporting Information.

## Acknowledgments

Armands Kazia acknowledges LIOS student grant (grant No IG-2026-06) and recovery and reseliance grant for PhD students (ANM_OSI_DG_09). The work was also supported by funding to Michael J. Blackman from the Wellcome Trust (220318/A/20/Z) and the Francis Crick Institute (https://www.crick.ac.uk/) which receives its core funding from Cancer Research UK (CC2129), the UK Medical Research Council (CC2129), and the Wellcome Trust (CC2129). For the purpose of Open Access, the author has applied a CC BY public copyright licence to any Author Accepted Manuscript version arising from this submission.

